# Girdin: an essential component of pre-replicative complex in human cells

**DOI:** 10.1101/634626

**Authors:** Lihong Wu, Yu Hua, Feiran Chang, Daochun Kong

## Abstract

A central event in the initiation of DNA replication in eukaryotes is the assembly of pre-replicative complex (pre-RC) on specific chromatin sites known as DNA replication origins. The pre-RC assembly process differs between budding and fission yeasts. In fission yeast, Sap1 directly participates in pre-RC assembly, together with the four initiation factors: ORC, Cdc18/Cdc6, Cdt1, and MCM. In metazoans, the nature of DNA replication origins is not defined and the mechanism of pre-RC assembly remains incompletely known. In this study, Girdin was identified as an essential replication initiation factor in human cells. Similar to the activity of Sap1, human Girdin binds to DNA origins, interacts with ORC, and is required for pre-RC assembly due to its essential role in recruitment of Cdc6 to DNA origins. Thus, DNA origins in human or metazoans are defined as including two elements, one bound by ORC and the other bound by Girdin.

## Introduction

In eukaryotes, the biochemical process of DNA replication initiation is divided into two steps. In the first step, a pre-replication complex (pre-RC) is assembled on specific chromatin sites or DNA replication origins at late M and G1 phases of the cell division cycle. In the second step, when cells are at the transition of G1 to S phase, the pre-RCs are activated by CDK and DDK kinases (1), triggering the recruitment of Cdc45 and GINS for the formation of CMG (Cdc45-MCM-GINS) replicative helicase, and the recruitment of other replication factors that act at replication forks, such as RPA, DNA pol α-primase, DNA pol δ, and DNA pol ε. Once these factors are assembled, DNA replication starts.

In budding yeast, pre-RC is assembled by four protein factors—ORC (origin recognition complex), Cdc6, Cdt1, and MCM (mini-chromosome maintenance). In the early 1970s, *Cdc6* gene was first identified by Hartwell and his colleagues in a screen of temperature-sensitive mutants defective in gene functions needed at specific stages of the cell division cycles (2). Their studies suggested that Cdc6 may specify a function necessary for the proper initiation of DNA replication, but its exact function was unknown at that time (3). Over the next twenty years, studies carried out in several labs finally revealed the direct participation of Cdc6 in pre-RC assembly, with Cdc6 essential for the recruitment of MCM to DNA origins (4–12). In a similar genetic screen, several temperature-sensitive mutants, named *cdc21, nda1*, and *nda4* in the fission yeast *S. pombe* and CDC54, CDC46, and CDC47 in the budding yeast *S. cerevisiae*, were isolated in the early 1980s (13–18). These mutants were later determined to harbor mutations in Mcm2, Mcm4, Mcm5, and Mcm7 gene, respectively. Just one or two years later, mutants defective in maintaining circular minichromosomes (Mcm^−^) were also isolated in *S. cerevisiae* (19,20). MCM was later identified to function in fungi to *homo sapiens* as a complex composed of six distinct and highly conserved subunits (21–27). ORC was first isolated in the budding yeast *S. cerevisiae* in 1992 (28). ORC is a six-subunit complex that remains bound to DNA origins throughout the cell cycle. Cdt1 was first identified in the fission yeast *S. pombe* (29). The *cdt1* gene is essential in fission yeast; cells with a null allele of *cdt1* arrest in the G1 phase, suggesting that Cdt1 function is related to DNA replication with either a direct role as a replication factor or an indirect regulator, such as a transcription factor that could promote entry into S phase (29). Cdt1 was subsequently found to be required for loading MCM to DNA origins for pre-RC assembly (30,31). Using recombinant ORC, Cdc6, Cdt1, and MCM, pre-RC could be reconstituted *in vitro* in the budding yeast *S. cerevisiae*, suggesting that these four proteins are sufficient for the assembly of pre-RC in budding yeast (32–34).

As defined, pre-RC assembles on DNA replication origins. However, the structures of DNA origins are remarkably different from budding yeast to fission yeast and metazoans. In the budding yeast *S. cerevisiae*, DNA origins are approximately 100 bp in size. Each origin contains a consensus sequence of 11 bp termed an “A” element that is essential for origin activity and is recognized and bound by ORC (35–38). In the fission yeast *S. pombe*, DNA origins range from 500-1500 bp in size (39–42), making them five to ten times larger than DNA origins in *S. cerevisiae*. The *S. pombe* DNA origins are large because they possess two essential elements that are recognized and bound by two origin recognition proteins, ORC and Sap1 (43,44). The average distance between ORC- and Sap1-bound elements ranges from ~300 to ~700 bp, and this distance combined with the size of the two origin elements explains the large size of DNA origins in *S. pombe*. Sap1, as an origin recognition factor, exhibits the same functions as ORC (44). Sap1 binds to DNA origins throughout the cell division cycle, Sap1 interacts with ORC to form a complex, and Sap1 is required for pre-RC assembly because it recruits Cdc18 (the homologue of Cdc6 in fission yeast) to DNA origins (44). Therefore, DNA origins in *S. cerevisiae* contain a single element that is bound by ORC, but those in *S. pombe* possess two discrete elements and two origin recognition factors, ORC and Sap1.

Similar to what occurs in budding and fission yeasts, the initiation of DNA replication in metazoans also occurs at specific chromosome sites, however, the nature of metazoan DNA origins remains elusive (45–54). DNA origins in metazoans are large (55–58), contain AT-rich sequences important for origin activity (42,59–61), and lack an easily recognizable consensus sequence. These three characteristics also generally describe *S. pombe* DNA origins, suggesting the possibility that metazoan and *S. pombe* DNA origins may have a similar structure. Initially, it was though that *S. pombe* DNA origins are AT-rich but lack a consensus sequence. Later, it was found that the *S. pombe* ORC binds to asymmetric AT-rich sequences with A in one strand and T in the other strand, in a process that depends on its 9-AT hook motifs (42,43,62,63). And Sap1 binds to a DNA sequence of 5’-(A/T)_n_(C/G)(A/T)_9–10_(G/C)(A/T)_*n*_-3’ (*n* ≧ 1) (44). Thus, based on the recent findings DNA origins in fission yeast do exhibit a certain level of sequence specificity. However, it remains to be determined if DNA origins in metazoans are similar to those in *S. cerevisiae* or *S. pombe*.

In this study, Girdin was identified in human cells as the homologue of the pre-RC component Sap1. Girdin is a highly conserved protein in metazoans, and sequence alignment indicates that *h*Girdin contains a region of ~200 amino acids that is highly homologous (~41% amino acid identity) to the highly conserved middle region of Sap1 protein in fission yeast. Sap1 has 254 amino acids in total. Like Sap1, *h*Girdin binds to DNA origins, physically interacts with ORC, and is required for loading Cdc6 to DNA origins for pre-RC assembly. The Sap1-homologous region in *h*Girdin can partially complement the function of Sap1 in fission yeast, further suggesting that Girdin is the homologue of Sap1 in metazoans. Thus, like those in fission yeast, DNA origins in human/metazoan cells possess two essential elements, with one bound by ORC and the other by Girdin. The assembly of pre-RC in metazoans requires the five components of ORC, Girdin, Cdc6, Cdt1, and MCM.

## Results

### Identification of the Sap1 homologue in metazoans

The similarity of DNA replication origins between fission yeast and metazoans suggests the likelihood of a Sap1 homologue in metazoans. Database-searching and immunoprecipitation against human Cdc6 identified the human protein Girdin. Girdin contains a region (~200 amino acids) that is highly homologous to Sap1 (Fig. 1*A, B*), with ~41% amino acids identity (with D/E, I/L, and R/K considered identical amino acids). The homologous region in Sap1 consists of amino acids 8 to 210, and this region is the most conserved in Sap1 proteins in fission yeast. Girdin is a very conserved protein present from *C. elegans* to human. Comparison of Girdin proteins from different organisms reveals that the Sap1-homologous region is the most conserved region, suggesting this region is probably the most important region for Girdin function. Girdin is expressed in all examined tissues and cell lines (64,65).

**Figure 1.**
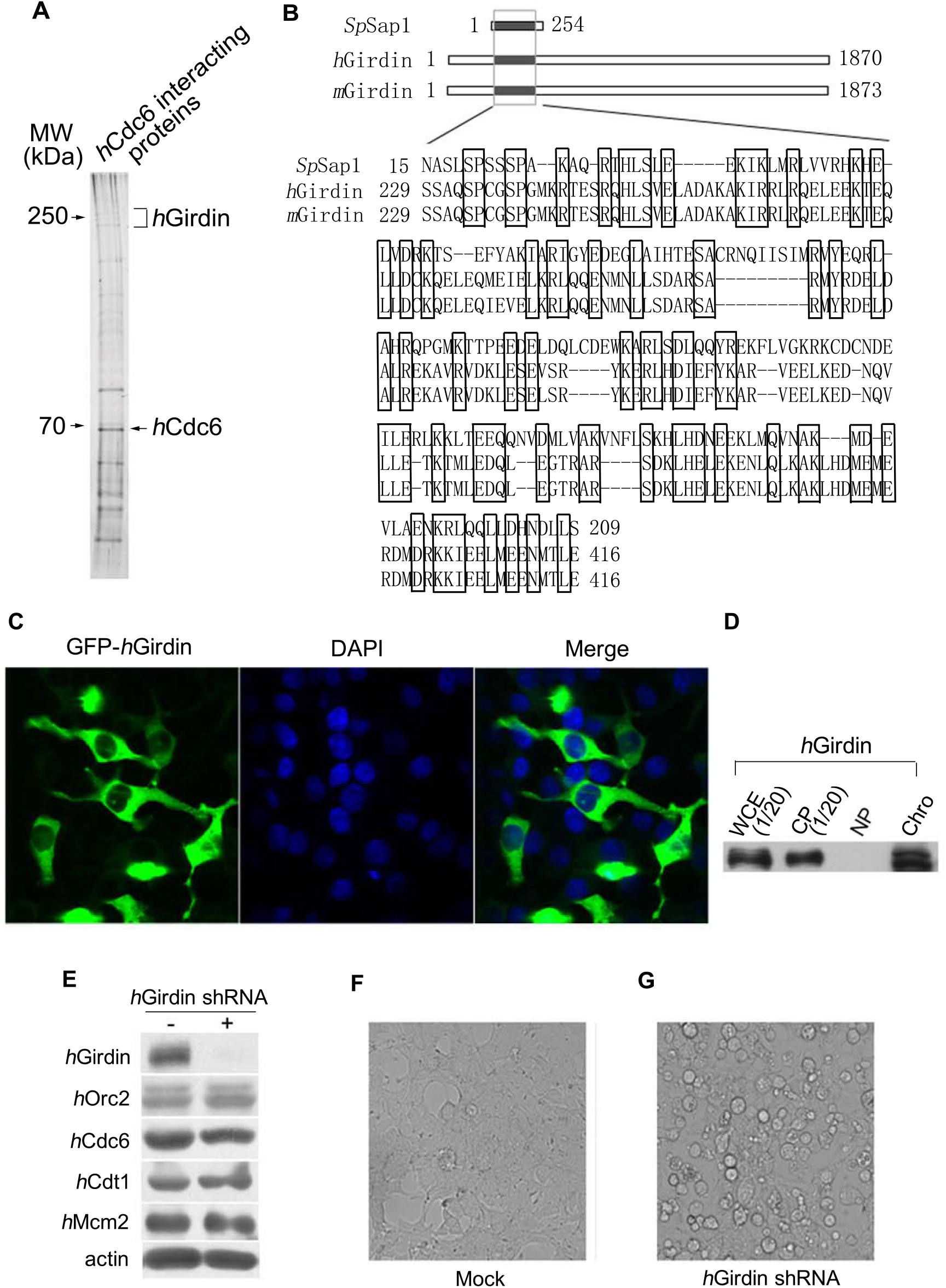
Girdin contains a region homologous to Sap1. *A*, Girdin interacts with Cdc6. A tagged version of Cdc6, 6His-3HA-Cdc6, was overexpressed in 293T cells and subsequently isolated by gel filtration, Ni^2+^-column, IP against HA, and glycerol gradient sedimentation. The silver-stained gel is presented and the protein bands of Cdc6 and *h*Girdin that were identified by mass spectroscopy are indicated. *B*, pairwise comparison of Sap1 and Girdin from human and mouse. *C* and *D*, the distribution of *h*Girdin in 293T cells. The majority (~90%) of *h*Girdin is localized in cytoplasm and the rest (~10%) is in the cell nucleus based on the fluorescent assay of GFP-*h*Girdin (*C*) and Western blotting of the cellular fractions (*D*). (*E-G*), Girdin is essential for cell growth. *E*, Girdin was depleted in 293T cells by Girdin shRNA. After nine days of incubation, the Girdin shRNA-treated cells were collected to measure the amounts of Girdin and the other pre-RC components in whole cells. Western blotting results are presented. (*F, G*), the Girdin shRNA-treated cells showed cell death (*G*) but the control shRNA-treated cells did not show death after ten days of incubation (*F*). Microscopic images of the cells are presented.

### Girdin is an essential protein for cell growth

To investigate whether Girdin, like Sap1, is essential in pre-RC assembly, we first examined its cellular distribution. Fluorescent analysis of GFP-tagged Girdin revealed that the majority of Girdin was in the cytoplasm, but a weak fluorescent signal was also detected in nuclei (Fig. 1*C*). To determine whether Girdin in the nucleus associates with chromatin, the fractions of cytoplasm, nucleoplasm, and chromatin were carefully separated and the amount of Girdin in each fraction was quantified by Western blotting. As shown in Fig. 1*D*, ~95% of Girdin was found in the cytoplasm and the remained (~5-10%) was in the nuclear fraction, where the Girdin was associated with chromatin.

Next, we examined whether Girdin is essential for cell growth. To do this, 293T cells were infected with lentiviral particles harboring either Girdin shRNA or control shRNA. After eight to nine days of incubation, the amount of Girdin was reduced by more than 90% in Girdin shRNA-introduced cells but the levels of the other initiation proteins, Orc2, Cdc6, Cdt1, and Mcm2, remained the same (Fig. 1*E*). At this point, Girdin shRNA-treated cells grew more slowly compared to the control cells. Beginning at the tenth or eleventh day, ~30-40% of Girdin shRNA-treated cells were dying or dead, but the control cells still grew well at the twentieth day or beyond (Fig. 1*F, G*). A similar result was obtained with HeLa cells or using a different shRNA. Efforts to knock out the gene of *h*Girdin by CRISPA-Cas9 were not successful. Together, these results indicate that Girdin is an essential protein for cell growth.

### The S phase progression of the cell cycle is affected when Girdin is reduced

Slower cell growth was observed in the Girdin siRNA- or shRNA-treated cells. To examine if Girdin affects the progression of S phase, the percentage of S phase cells was measured in the Girdin siRNA- or control siRNA-treated cells by BrdU incorporation and subsequent florescent assay and FACS analysis. Based on the percentage of cells that incorporated BrdU, ~53% of the Girdin siRNA-transfected cells were in S phase when Girdin was reduced by ~90%, while only ~41% of the control cells were in the S phase (Fig. 2*A, B*). The FACS analysis also clearly shows more cells in the S phase when Girdin was reduced (Fig. 2*C*). This result is consistent with that of a previous study showing that the depletion of another replication initiation protein Treslin from human cells increases the number of cells in S phase (66). Consistent with slower progression of S phase, DNA synthesis was inhibited by ~20-35% in the Girdin siRNA-treated cells compared to the control cells (Fig. 2*D*).

**Figure 2.**
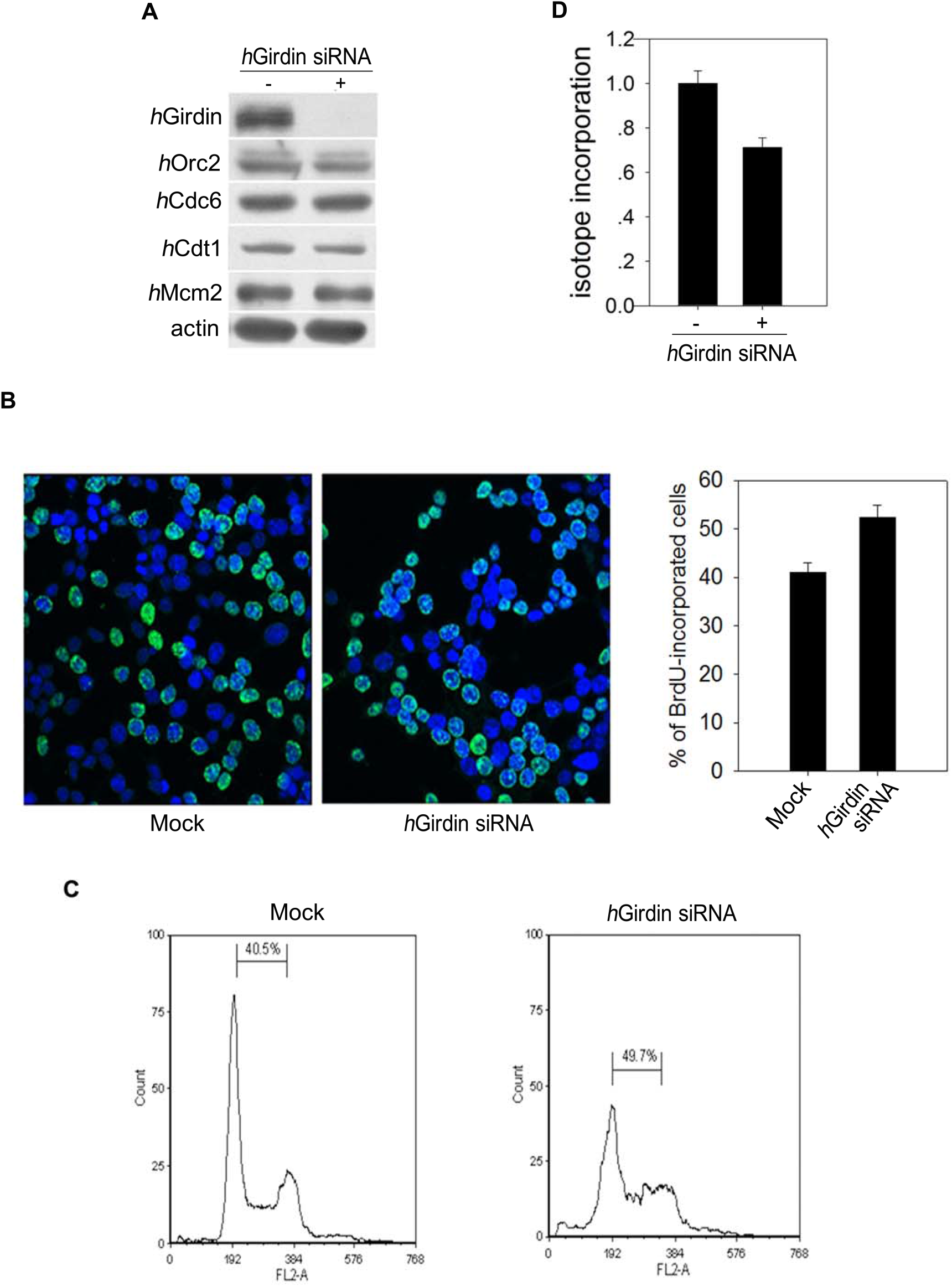
Girdin is required for the S phase progression in cell division cycles. 293T cells were transfected with Girdin siRNA or control siRNA and then incubated for three days before the assays (*A-C*) were carried out. *A*, cellular amounts of Girdin and the other pre-RC components in the Girdin siRNA- or control siRNA-treated cells, as determined by Western blotting. *B*, slow S phase progression in the Girdin siRNA-treated cells. Mock- or Girdin siRNA-treated 293T cells were labeled for 30 min in medium containing 200 μg/ml of BrdU. The incorporated BrdU was detected with mouse anti-BrdU antibody and FITC-labeled goat antibody against mouse IgG. The percentage of cells showing BrdU incorporation in normally growing cells (Mock) or *h*Girdin siRNA-treated cells was statistically measured and is presented in the right panel. *C*, FACS analysis of the Girdin siRNA- or control siRNA-treated 293T cells. *D*, the inhibition of DNA synthesis in *h*Girdin siRNA-treated cells. After incubation of the *h*Girdin siRNA- or control siRNA-treated 293T cells for 20 hrs, nocodazole was added to the cultures for 15 hrs to arrest cells at G2/M phase in the presence of siRNA. Then, the cells were released into S phase with fresh medium with additional incubation of 9.5 to 11.5 hrs. FACS analysis indicated that the *h*Girdin siRNA- and control siRNA-treated 293T cells exhibited the same rate to progress into S phase from the G2/M arrest in a parallel test (data not shown). [α-^32^P] dATP and 0.008% digitonin (final concentration) were added to medium. After DNA synthesis with isotope incorporation was allowed for 30 min, cells were then collected and genomic DNA was purified. To measure the rate of DNA synthesis, the incorporated isotope was measured for the same amount of genomic DNA from *h*Girdin siRNA- or control siRNA-treated cells.

### Girdin and ORC bind to the same regions on genomic DNA and they physically interact with each other

To determine whether Girdin and ORC bind to the same regions on genomic DNA, their genomic binding sites were identified with ChIP-seq assays. As shown in Fig. 3*A*, the genomic binding sites of Girdin and ORC are the same. The completely overlapping binding sites suggests that Girdin and ORC are under physical interaction on chromatin DNA. To confirm that, reciprocal IPs were performed. The results in Fig. 3*B* show that IP against Girdin or ORC brought down ORC (Orc2 subunit) or Girdin, respectively, indicating that like Sap1 and ORC in *S. pombe*, human Girdin and ORC physically interact with each other. The association of Girdin with DNA was also examined during the cell cycle. The result shown in Fig. 3*C* indicates that Girdin, like ORC, binds to DNA origins throughout the cell cycle.

**Figure 3.**
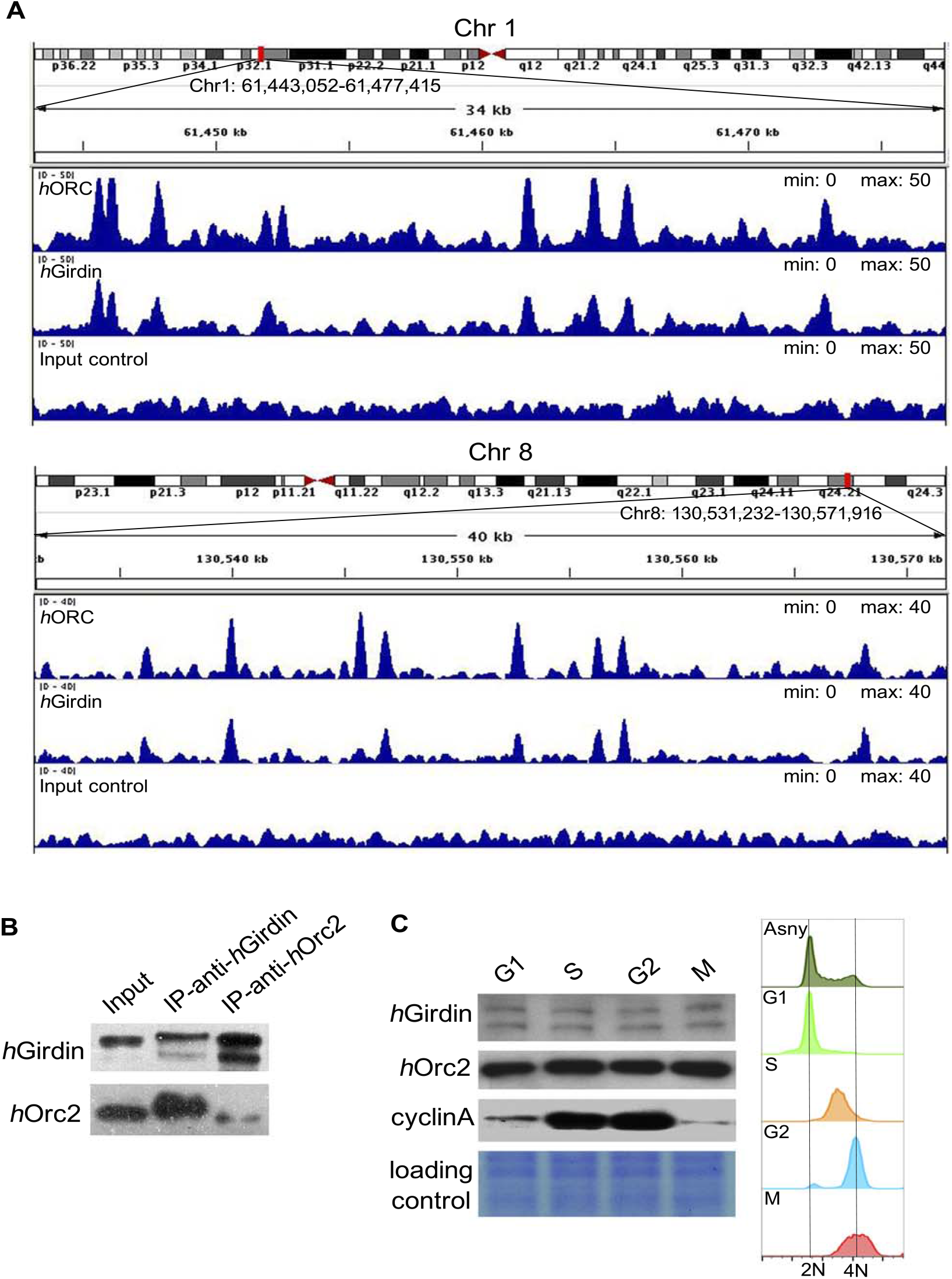
*h*Girdin and ORC bind to the same DNA regions/origins on human genomic DNA. *A*, ChIP-seq against Orc2 subunit or *h*Girdin was carried out as described in methods. Presented are examples showing the overlapped *h*Girdin and ORC binding sites in two regions from Chromosome #1 and #8. *B*, reciprocal IPs reveal the interaction of Girdin and ORC. Presented is the result of Western blotting assay. C, Girdin or ORC (Orc2) present at a constant level on chromatin DNA throughout the cell division cycle. 293T cells at ~25% confluency were first treated with 2 mM thymidine at 37°C for 20 h, then released to fresh medium without drugs for 8 h. Then, 20 ng/μL of nocodazole was added to the culture for 12 hrs to arrest cells in G2/M phase. The cells were released to fresh medium and cell samples were taken at 0 h (M phase), 4.5 h (G1 phase), 9 h (S phase), and 16 h (G2 phase) for FACS analysis and Western blotting assay. The levels of *h*Girdin, *h*Orc2, and cyclin A were quantified by Western blotting assays by using the corresponding antibodies (left panel). The right panel shows the results of FACS analyses. The cell phases are indicated: Asny (dark green), G1 (light green), S (orange), G2 (blue), and M (red).

### Girdin is required to recruit and load Cdc6 onto DNA for pre-RC assembly

The interaction between Girdin and Cdc6 was further confirmed by performing reciprocal IPs with cell extracts containing a physiological level of Cdc6 (Fig. 4*A*). Next, we examined the requirement of Girdin for Cdc6 loading onto DNA for pre-RC assembly. ORC- and MCM-dependent *S. cerevisiae* or human cell-free systems for pre-RC assembly and DNA replication were previously reported (32–34,67–69). In this assay, nuclear extracts prepared from 293T cells in late G1 phase were used. During IP depletion experiments, we identified ORC, Girdin, Cdc6, Cdt1, and MCM in sub-complexes even in nuclear extracts free of genomic DNA. Thus, immuno-depletion of Girdin or Orc2 was carried out in the presence of 0.6 M KCl. For removal of ~95-98% of ORC or Girdin from the extracts, immuno-depletion of Girdin or Orc2 was routinely conducted two or three times. As shown in Fig. 4*B*, this method allowed effective removal of Girdin or Orc2 subunit, but not the other pre-RC components. Assembly of pre-RC was performed on salmon sperm DNA at a salt concentration of 0.1 to 0.15 M (see Methods). The result presented in Fig. 4*C* shows that Girdin was required to load Cdc6 and, subsequently, MCM to DNA, but was not required for Cdt1 loading. Additionally, the binding of ORC to DNA was not affected in the absence of Girdin, and that ORC, as shown previously, was required to load Cdc6 and Cdt1 and then MCM to DNA. In a control reaction, Cdc6, Cdt1, and MCM were loaded onto DNA when both Girdin and ORC were present. To verify that the absence of Girdin was responsible for the observed disruption of the pre-RC assembly (Fig. 4*C*), recombinant Girdin, which was overexpressed and subsequently purified from *S. pombe* cells (Fig. 4*D*), was added back to the Girdin-depleted extract and then the assembly of pre-RC was restored (Fig. 4 *E*).

**Figure 4.**
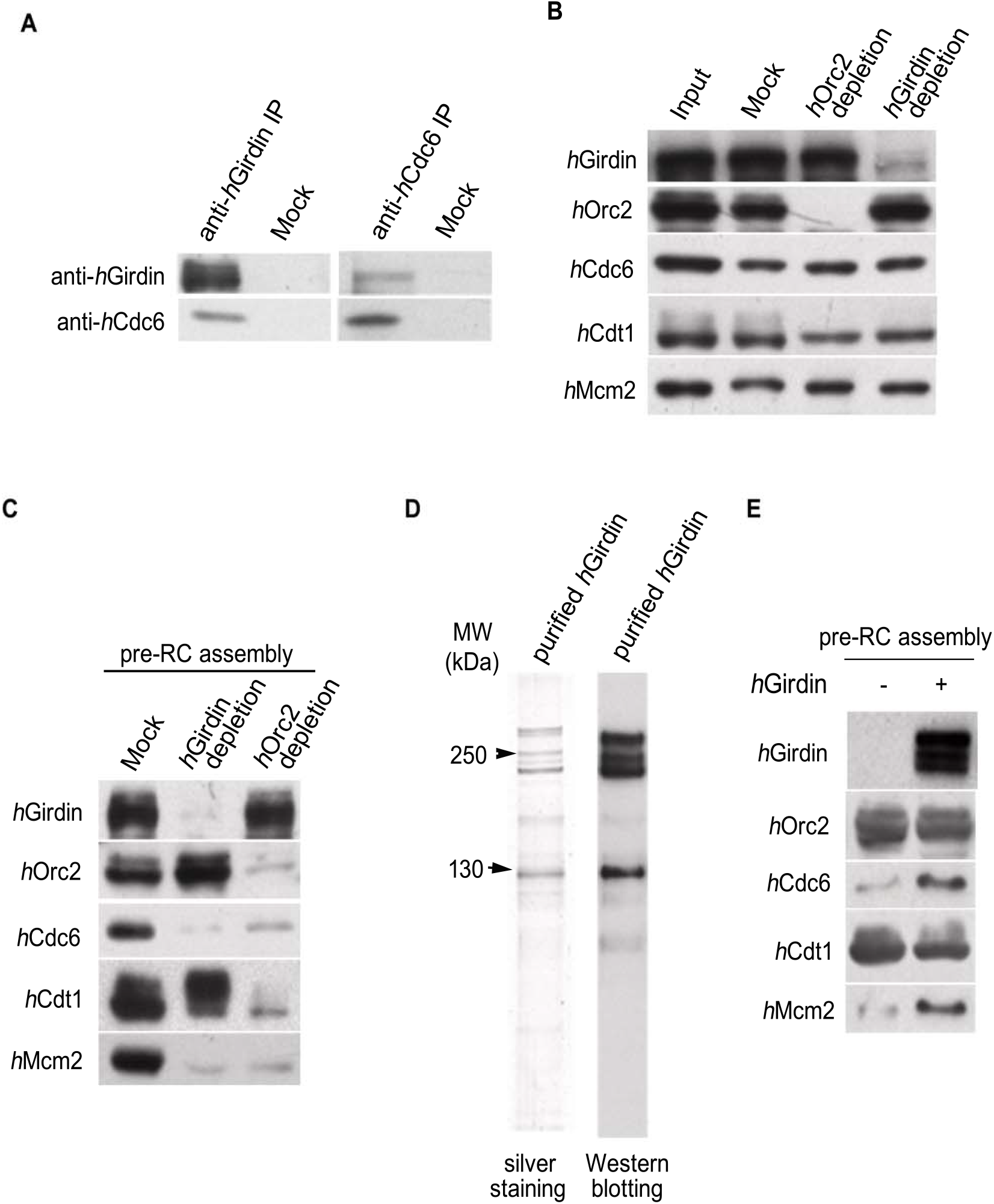
*h*Girdin is required to load Cdc6 to DNA for pre-RC assembly. *A*, the reciprocal IP shows that *h*Girdin interacts with Cdc6. This assay was carried out with cell extracts prepared from cells expressing a physiological level of *h*Girdin or Cdc6. *B-E*, Girdin is required to load Cdc6 to DNA for pre-RC assembly. *B*, the immunodepletion of Girdin or Orc2 from the nuclear extracts of 293T cells in late G1 phase. *C*, the assembly of pre-RCs on Salmon sperm DNA-cellulose with *h*Girdin-, or Orc2-, or mock-depleted nuclear extracts. *D*, recombinant Girdin was overexpressed and subsequently purified from *S. pombe* cells. *E*, the addition of recombinant Girdin to the Girdin-depleted extract restores the loading of Cdc6 and MCM onto DNA. Presented above are the Western blotting results except the silver-stained *h*Girdin purified from *S. pombe* cells.

### Girdin complements the function of Sap1 in fission yeast *S. pombe*

To confirm that Girdin is functionally related to Sap1, we asked if Girdin could complement the function of Sap1 in the fission yeast. A plasmid that expresses the Sap1-homologous region of Girdin under the control of nmt41 promoter was transformed into the fission yeast *Sp-dk1-sap1*^ts5^ cells and the transformed cells were examined for their ability to grow at the restrictive temperature of 32°C or 34°C. The results in Fig. 5*A, B* show that the expression of the Sap1-homologous region of Girdin could enhance the survival rate of *Sp-dk1-sap1*^ts5^ cells at 32°C or 34°C by ~5 to 25 folds. This result indicates that the Sap1-homologous region of Girdin can partially complement the function of Sap1 in the fission yeast, consistent with the conservation of biological function from Sap1 to Girdin.

**Figure 5.**
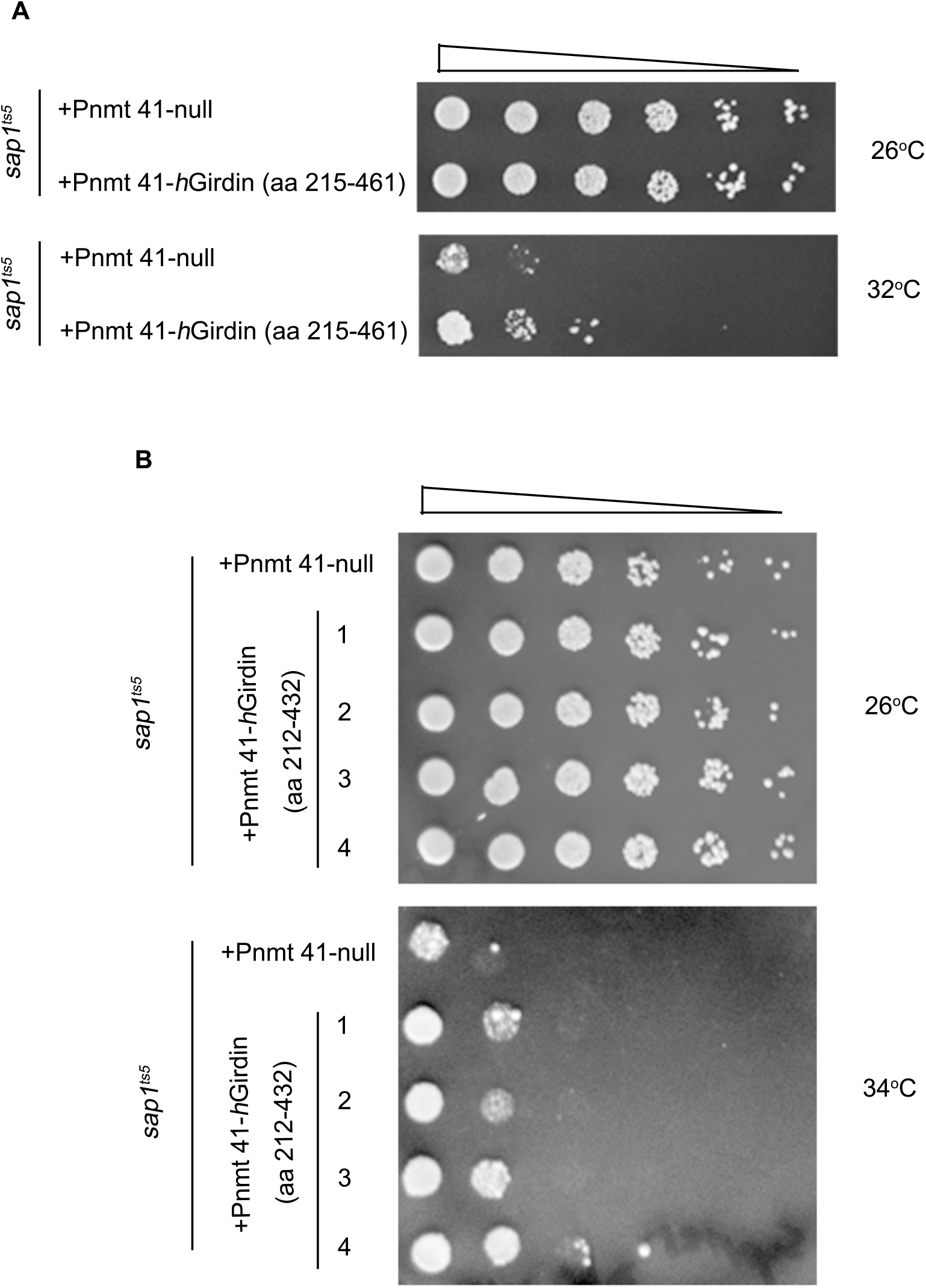
The Sap1-homologous region in Girdin partially complements the function of Sap1 in *Sp*-dk1-*sap1*^ts5^ cells. A recombinant plasmid expressing the indicated portion of *h*Girdin sequence was constructed and transformed into *Sp*-dk1-*sap1*^ts5^ strain. A five-fold dilution test was carried out to examine the survival rate of the corresponding strains at 32°C or 34°C. The nmt-41 promoter is a middle strength promoter from which gene expression is induced by the absence of thiamine. A, the Sap1-homologous region (aa 215-461) in *h*Girdin partially complements the function of Sap1 in *Sp-dk1-sap1*^ts5^ at 32°C. *B*, the Sap1-homologous region (aa 212-432) in *h*Girdin partially complements the function of Sap1 in *Sp-dk1-sap1*^ts5^ at 34°C. Four independent clones were examined.

**Figure 6.**
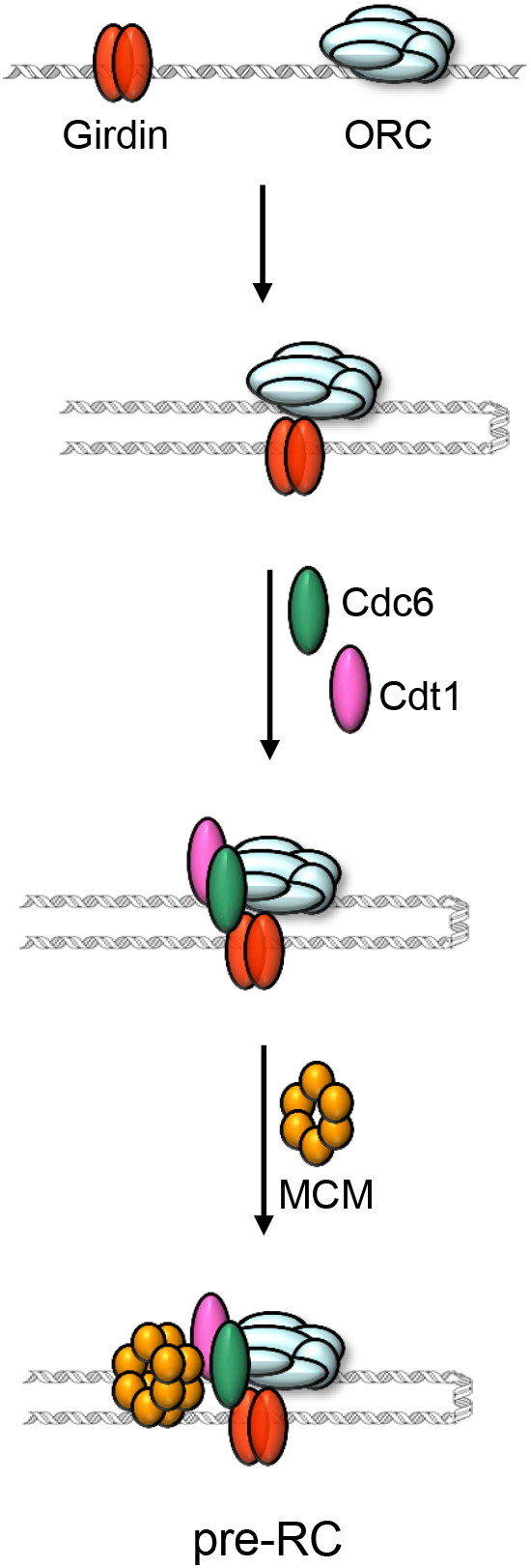
Schematic of pre-RC assembly in human cells. Girdin and ORC bind to specific DNA sequences, or DNA replication origins, on human genomic DNA. They form a complex through physical interaction and then stay associated with genomic DNA throughout cell division cycle. In late M and G1 phases of the cell division cycles, the Girdin-ORC complex loads Cdc6 and Cdt1 onto replication origins, and Cdc6 and Cdt1 then recruit MCM complex to complete pre-RC assembly. Pre-RCs are subsequently activated at the transition of G1 to S phase. After recruitment of other replication proteins to DNA origins, DNA replication finally starts.

## Discussion

There is significant experimental evidence that DNA replication initiates at specific sites in chromosomes in metazoans. However, the molecular mechanism for metazoan cells to choose replication initiation sites remains unclear. In particular, the nature of metazoan DNA replication origins and how DNA replication is initiated by assembly of pre-RC have been unresolved for more than two decades, significantly hampering further elucidation of the mechanism of replication initiation in metazoans.

The identification of a second DNA origin recognition factor, Sap1, in *S. pombe* suggests that a Sap1 homologue may also exist in metazoans, since *S. pombe* and metazoans appear to have a similar origin structure. By searching for proteins that interact with Cdc6 in human cells, we identified Girdin, which contains a region highly homologous to a region of ~200 amino acids in Sap1. This homologous region is also highly conserved among Sap1 proteins in the fission yeast species whose genomic sequences are available. However, the N-terminal ~10 amino acids and C-terminal ~40-50 amino acids in Sap1 are not very conserved. The nearly 41% identity of amino acids in the homologous region between Sap1 and *h*Girdin suggests that Girdin is a homologue of Sap1 in metazoans, and this was subsequently demonstrated by several lines of experimental evidence. First, like Sap1 and other replication proteins acting either in replication initiation or at replication forks, a reduced level of *h*Girdin slowed S phase progression (Fig. 2*A-D*). Second, like Sap1, *h*Girdin binds to DNA regions that are bound by ORC, and *h*Girdin and ORC physically interact as a complex on DNA origins. This *h*Girdin-ORC complex provides the biochemical basis for their collective action in recruiting Cdc6 to replication origins (Fig. 3*A, B*). Third, like Sap1, *h*Girdin is required to recruit Cdc6 to DNA origins (Fig. 4). Finally, a functional complementation test revealed that the homologous region in *h*Girdin partially complemented Sap1 function (Fig. 5), suggesting that *h*Girdin possesses a function that Sap1 also utilizes. Taken together, the experimental evidence presented in this study indicates that *h*Girdin is a genuine homologue of Sap1 and functions in pre-RC assembly and replication initiation.

Why do human/metazoan cells, like the fission yeast *S. pombe*, require a factor in addition to ORC for their pre-RC assembly? The answer to this question is likely related to the nature of the ORC complex. There are some obvious differences among ORC complexes in the budding yeast *S. cerevisiae*, the fission yeast *S. pombe*, and metazoans. First, the Orc6 subunits (~28-30 kDa) in fission yeast and metazoans differ from *S. cerevisiae* Orc6 (~50 kDa) in size and amino acid sequence. Second, *S. pombe* Orc4 contains an additional domain (~60 kDa) that contains 9 AT-hook motifs at its N-terminus. This AT-hook domain is not present in *S. cerevisiae* ORC and is solely responsible for binding of *S. pombe* ORC to AT-rich sequences in DNA origins(42,43,62,63). Third, a mutant *S. pombe* ORC that lacks the 9 AT-hook domain shows random, low affinity binding to DNA (62). Fourth, human ORC, similar to *S. pombe* ORC lacking the 9 AT-hook motifs, appears to have a sequence-independent DNA binding property (70). Fifth, *S. cerevisiae* ORC interacts with DNA using its five subunits (71), which is significantly different from *S. pombe* ORC which uses only one domain in Orc4 (the 9 AT-hook motifs) to interact with DNA. Whether human or metazoan ORC utilizes an accessory subunit similar to the 9 AT-hook motifs in *S. pombe* ORC remains to be determined. It is possible that such a subunit exists and promotes human/metazoan ORC binding to DNA. The difference in ORC and ORC-DNA interaction may explain the requirement of Sap1/Girdin for pre-RC assembly in fission yeast and metazoans.

The identification of replication initiation factor Girdin provides insight into the structure of DNA origins in metazoans. Like DNA origins in fission yeast, DNA origins in metazoans possess two origin elements, one that is recognized and bound by ORC and one that is recognized and bound by Girdin. Thus, metazoan DNA origins are defined as possessing two discrete elements that are separately bound by two origin recognition factors. Future investigation of metazoan DNA origins should determine how Girdin interacts with DNA and whether metazoan ORC associates with an accessory factor that is analogous to the AT-hooks domain in *S. pombe* Orc4 subunit.

## Experimental procedures

### Purification of *h*Girdin and preparation of polyclonal antibodies against human Orc2, Mcm2, Cdc6, Cdt1, and the Sap1-homologous region of *h*Girdin

The tagged *h*Girdin was overexpressed as 6His-*h*Girdin in *S. pombe* cells. All expressed 6His-Girdin was in the nuclei and the majority bound to chromatin. The 6His-Girdin in the chromatin extract was first precipitated by 40% ammonium sulfate. The pellet containing 6His-Girdin was then resuspended in buffer A (50 mM HEPES-HCl [pH 7.5], 100 mM KCl, 5 mM magnesium acetate, 5 mM dithiothreitol, 1 mM EDTA, 1 mM EGTA, 10% glycerol, 0.04% NP-40, and protease inhibitors (Sigma)) and applied to Sephacryl S-300 column for filtration chromatography. The fractions containing 6His-Girdin were then applied to Q-sepharose column and 6His-Girdin was eluted at 0.5 to 0.6 M KCl. Then, the eluted 6His-Girdin was further purified with Ni^2+^-column and eluted at 250 mM imidazole. The purified 6His-Girdin was dialyzed against buffer A and then stored in −80°C freezer.

A polyclonal antibody against the Sap1-homologous region of *h*Girdin was obtained by immunizing rabbits with recombinant protein of the Sap1-homologous region of Girdin that was overexpressed in *E. coli* and purified. Polyclonal antibodies against human Orc2, Mcm2, Cdc6, Cdt1 were similarly obtained.

### ChIP-seq assays to determine the binding sites of human ORC and *h*Girdin on genomic DNA

To determine human ORC and *h*Girdin binding sites, ChIP-seq assay was carried out as described (Euskirchen et al., Genome Res 17, 898-909; Johnson et al., Science 316, 1497-1502) with some modifications. First, 293T cells were grown to the confluence of 70% in DMEM containing 10% fetal bovine serum, and then arrested at G2/M phase after treatment of the culture with nocodazole (50 μg/ml) for 12 hrs. The chromatin-bound proteins were cross-linked to DNA by treating the cells with 1% formaldehyde for 10 min. The formaldehyde-treated cells were collected, washed with PBS twice, and then homogenized to obtain nuclei. The nuclei were lysed with RIPA Buffer (1XPBS / 1% NP-40 / 0.5% sodium deoxycholate / 0.1% SDS) and the chromatin was sheared to an average of 250 bp with sonication. The ChIP was then carried out to obtain ORC or Girdin-bound chromatin fragments using polyclonal antibody against Orc2 or *h*Girdin.

### RNA interference

In total, five sets of siRNA sequences were tested. Three of them effectively mediated silencing of Girdin expression. One sequence is 5’-GCAATTAGAGAGTGAACTA-3’ (nucleotide 2340-2358 in *h*Girdin cDNA gene, only sense sequence is shown). This sequence was purchased as a 21 nucleotide synthetic duplex. 293T cells were transfected with the siRNA or a 21-nucleotide non-related RNA as a control by using lipofectamine 2000 (Invitrogen) according to the manufacturer’s protocol. For shRNA-mediated knockdown of *h*Girdin, the targeted sequence is 5’-GAAGGAGAGGCAACTGGAT-3’ (nucleotides 4166-4184, only the sense sequence is shown). A set of single-stranded oligonucleotides encoding the Girdin target shRNA and its complement was synthesized, annealed, and inserted into lentiviral shRNA expression vector pLKO.1 (forward oligo, 5’-CCGGGAAGGAGAGGCAACTGGATCTCGAGATCCAGTTGCCTCTCCTTCTTTTTG-3’; reverse oligo, 5’-AATTCAAAAAGAAGGAGAGGCAACTGGATCTCGAGATCCAGTTGCCTCTCCTTC-3’). The production of lentiviral particles and lentiviral infection were performed according to manufacturer’s instructions.

### Preparation of nuclear extract from 293T cells at late G1 phase and immune-depletion of Girdin or Orc2

The 293T cells were incubated in DMEM medium containing 10% fetal bovine serum, were first arrested at G1/S phase in the presence of thymidine (2 mM) for ~20 hrs, and were then released to G2/M phase in the presence of nocodazole (50 μg/ml) for 14 hrs. The cells were then released from nocodazole arrest and grew into late G1 phase by incubation in fresh medium for 6.5 hrs. The nuclei obtained with hypotonic buffer treatment and homogenization were frozen once and then crushed at high speed (37000g) to obtain the nucleoplasm supernatant. The pellet containing chromatin was sequentially extracted with buffer A containing 0.3 and then buffer A containing 0.6 M KCl. These extracts were mixed with the nucleoplasm to obtain whole nuclear extract at a salt concentration of ~0.2 M. To specifically remove ORC or Girdin protein from the nuclear extract, the salt concentration in the nuclear extract was first adjusted to 0.6 M to prevent removal of ORC or Girdin-interacting proteins such as Cdc6 during immunodepletion. The immunodepletion of Orc2 or Girdin was performed two or three times, allowing removal of more than 95% to 98% of Orc2 or Girdin from the extract. As a positive control in the subsequent pre-RC assembly reaction, whole nuclear extract samples were also mock-depleted with rabbit IgG.

### Assembly of pre-RC on salmon sperm DNA-cellulose with human 293T nuclear extracts

The mock-, Orc2-, or *h*Girdin-depleted nuclear extracts were dialyzed against buffer A to adjust the salt concentration to 0.15 M. In a standard pre-RC assembly reaction, ~0.5 ml of nuclear extract from 293T cells at late G1 phase was mixed with 50 μl of DNA-cellulose in the presence of 5 mM ATP, 3 mM DTT, 10 mM of creatine phosphate, 20 units of creatine phosphokinase/ml, and proteinase inhibitors. The reactions were incubated at 4°C for 30 min, and then spun and the supernatant was removed. The pellet was washed with one ml of buffer A four times and then washed quickly twice with 0.5 ml of buffer A with salt (0.3 M KCl) to remove any proteins that were non-specifically bound to the salmon sperm DNA or cellulose. The pre-RC components assembled onto the DNA-cellulose were examined by SDS-PAGE electrophoresis and Western blotting assay.

## Acknowledgements

We thank the members of the Kong Lab for their support during the course of this project. This work was supported by grants from the Ministry of Science and Technology of China (2013CB911000 and 2016YFA0500301), the National Natural Science Foundation of China (no. 31661143032, no. 31230021, and no. 31730022), the Peking-Tsinghua Center for Life Sciences, and the National Key Laboratory of Protein and Plant Gene Research.

## References

1. Bell, S. P., and Dutta, A. (2002) DNA replication in eukaryotic cells. Annual review of biochemistry 71, 333–374

2. Hartwell, L. H., Mortimer, R. K., Culotti, J., and Culotti, M. (1973) Genetic Control of the Cell Division Cycle in Yeast: V. Genetic Analysis of cdc Mutants. Genetics 74, 267–286

3. Hartwell, L. H. (1976) Sequential function of gene products relative to DNA synthesis in the yeast cell cycle. Journal of molecular biology 104, 803–817

4. Lisziewicz, J., Godany, A., Agoston, D. V., and Kuntzel, H. (1988) Cloning and characterization of the Saccharomyces cerevisiae CDC6 gene. Nucleic acids research 16, 11507–11520

5. Zhou, C., Huang, S. H., and Jong, A. Y. (1989) Molecular cloning of Saccharomyces cerevisiae CDC6 gene. Isolation, identification, and sequence analysis. The Journal of biological chemistry 264, 9022–9029

6. Zhou, C., and Jong, A. (1990) CDC6 mRNA fluctuates periodically in the yeast cell cycle. The Journal of biological chemistry 265, 19904–19909

7. Palmer, R. E., Hogan, E., and Koshland, D. (1990) Mitotic transmission of artificial chromosomes in cdc mutants of the yeast, Saccharomyces cerevisiae. Genetics 125, 763–774

8. Hogan, E., and Koshland, D. (1992) Addition of extra origins of replication to a minichromosome suppresses its mitotic loss in cdc6 and cdc14 mutants of Saccharomyces cerevisiae. Proceedings of the National Academy of Sciences of the United States of America 89, 3098–3102

9. Liang, C., Weinreich, M., and Stillman, B. (1995) ORC and Cdc6p interact and determine the frequency of initiation of DNA replication in the genome. Cell 81, 667–676

10. Cocker, J. H., Piatti, S., Santocanale, C., Nasmyth, K., and Diffley, J. F. (1996) An essential role for the Cdc6 protein in forming the pre-replicative complexes of budding yeast. Nature 379, 180–182

11. Tanaka, T., Knapp, D., and Nasmyth, K. (1997) Loading of an Mcm protein onto DNA replication origins is regulated by Cdc6p and CDKs. Cell 90, 649–660

12. Aparicio, O. M., Weinstein, D. M., and Bell, S. P. (1997) Components and dynamics of DNA replication complexes in S. cerevisiae: redistribution of MCM proteins and Cdc45p during S phase. Cell 91, 59–69

13. Nasmyth, K., and Nurse, P. (1981) Cell division cycle mutants altered in DNA replication and mitosis in the fission yeast Schizosaccharomyces pombe. Molecular & general genetics: MGG 182, 119–124

14. Toda, T., Umesono, K., Hirata, A., and Yanagida, M. (1983) Cold-sensitive nuclear division arrest mutants of the fission yeast Schizosaccharomyces pombe. Journal of molecular biology 168, 251–270

15. Miyake, S., Okishio, N., Samejima, I., Hiraoka, Y., Toda, T., Saitoh, I., and Yanagida, M. (1993) Fission yeast genes nda1+ and nda4+, mutations of which lead to S-phase block, chromatin alteration and Ca2+ suppression, are members of the CDC46/MCM2 family. Molecular biology of the cell 4, 1003–1015

16. Coxon, A., Maundrell, K., and Kearsey, S. E. (1992) Fission yeast cdc21+ belongs to a family of proteins involved in an early step of chromosome replication. Nucleic acids research 20, 5571–5577

17. Moir, D., Stewart, S. E., Osmond, B. C., and Botstein, D. (1982) Cold-sensitive cell-division-cycle mutants of yeast: isolation, properties, and pseudoreversion studies. Genetics 100, 547–563

18. Hennessy, K. M., Lee, A., Chen, E., and Botstein, D. (1991) A group of interacting yeast DNA replication genes. Genes & development 5, 958–969

19. Maine, G. T., Sinha, P., and Tye, B. K. (1984) Mutants of S. cerevisiae defective in the maintenance of minichromosomes. Genetics 106, 365–385

20. Yan, H., Gibson, S., and Tye, B. K. (1991) Mcm2 and Mcm3, two proteins important for ARS activity, are related in structure and function. Genes & development 5, 944–957

21. Adachi, Y., Usukura, J., and Yanagida, M. (1997) A globular complex formation by Nda1 and the other five members of the MCM protein family in fission yeast. Genes to cells: devoted to molecular & cellular mechanisms 2, 467–479

22. Lei, M., Kawasaki, Y., and Tye, B. K. (1996) Physical interactions among Mcm proteins and effects of Mcm dosage on DNA replication in Saccharomyces cerevisiae. Molecular and cellular biology 16, 5081–5090

23. Dalton, S., and Hopwood, B. (1997) Characterization of Cdc47p-minichromosome maintenance complexes in Saccharomyces cerevisiae: identification of Cdc45p as a subunit. Molecular and cellular biology 17, 5867–5875

24. Su, T. T., Feger, G., and O’Farrell, P. H. (1996) Drosophila MCM protein complexes. Molecular biology of the cell 7, 319–329

25. Richter, A., and Knippers, R. (1997) High-molecular-mass complexes of human minichromosome-maintenance proteins in mitotic cells. European journal of biochemistry 247, 136–141

26. Romanowski, P., Madine, M. A., and Laskey, R. A. (1996) XMCM7, a novel member of the Xenopus MCM family, interacts with XMCM3 and colocalizes with it throughout replication. Proceedings of the National Academy of Sciences of the United States of America 93, 10189–10194

27. Thommes, P., Kubota, Y., Takisawa, H., and Blow, J. J. (1997) The RLF-M component of the replication licensing system forms complexes containing all six MCM/P1 polypeptides. The EMBO journal 16, 3312–3319

28. Bell, S. P., and Stillman, B. (1992) ATP-dependent recognition of eukaryotic origins of DNA replication by a multiprotein complex. Nature 357, 128–134

29. Hofmann, J. F., and Beach, D. (1994) cdt1 is an essential target of the Cdc10/Sct1 transcription factor: requirement for DNA replication and inhibition of mitosis. The EMBO journal 13, 425–434

30. Nishitani, H., Lygerou, Z., Nishimoto, T., and Nurse, P. (2000) The Cdt1 protein is required to license DNA for replication in fission yeast. Nature 404, 625–628

31. Maiorano, D., Moreau, J., and Mechali, M. (2000) XCDT1 is required for the assembly of pre-replicative complexes in Xenopus laevis. Nature 404, 622–625

32. Remus, D., Beuron, F., Tolun, G., Griffith, J. D., Morris, E. P., and Diffley, J. F. (2009) Concerted loading of Mcm2-7 double hexamers around DNA during DNA replication origin licensing. Cell 139, 719–730

33. Evrin, C., Clarke, P., Zech, J., Lurz, R., Sun, J., Uhle, S., Li, H., Stillman, B., and Speck, C. (2009) A double-hexameric MCM2-7 complex is loaded onto origin DNA during licensing of eukaryotic DNA replication. Proceedings of the National Academy of Sciences of the United States of America 106, 20240–20245

34. Tsakraklides, V., and Bell, S. P. (2010) Dynamics of pre-replicative complex assembly. The Journal of biological chemistry 285, 9437–9443

35. Palzkill, T. G., and Newlon, C. S. (1988) A yeast replication origin consists of multiple copies of a small conserved sequence. Cell 53, 441–450

36. Huberman, J. A., Zhu, J. G., Davis, L. R., and Newlon, C. S. (1988) Close association of a DNA replication origin and an ARS element on chromosome III of the yeast, Saccharomyces cerevisiae. Nucleic acids research 16, 6373–6384

37. Kearsey, S. (1983) Analysis of sequences conferring autonomous replication in baker’s yeast. The EMBO journal 2, 1571–1575

38. Marahrens, Y., and Stillman, B. (1992) A yeast chromosomal origin of DNA replication defined by multiple functional elements. Science 255, 817–823

39. Zhu, J., Brun, C., Kurooka, H., Yanagida, M., and Huberman, J. A. (1992) Identification and characterization of a complex chromosomal replication origin in Schizosaccharomyces pombe. Chromosoma 102, S7–16

40. Zhu, J., Carlson, D. L., Dubey, D. D., Sharma, K., and Huberman, J. A. (1994) Comparison of the two major ARS elements of the ura4 replication origin region with other ARS elements in the fission yeast, Schizosaccharomyces pombe. Chromosoma 103, 414–422

41. Clyne, R. K., and Kelly, T. J. (1995) Genetic analysis of an ARS element from the fission yeast Schizosaccharomyces pombe. The EMBO journal 14, 6348–6357

42. Kong, D., and DePamphilis, M. L. (2001) Site-specific DNA binding of the Schizosaccharomyces pombe origin recognition complex is determined by the Orc4 subunit. Molecular and cellular biology 21, 8095–8103

43. Kong, D., and DePamphilis, M. L. (2002) Site-specific ORC binding, pre-replication complex assembly and DNA synthesis at Schizosaccharomyces pombe replication origins. The EMBO journal 21, 5567–5576

44. Guan, L., He, P., Yang, F., Zhang, Y., Hu, Y., Ding, J., Hua, Y., Zhang, Y., Ye, Q., Hu, J., Wang, T., Jin, C., and Kong, D. (2017) Sap1 is a replication-initiation factor essential for the assembly of pre-replicative complex in the fission yeast Schizosaccharomyces pombe. The Journal of biological chemistry 292, 6056–6075

45. Handeli, S., Klar, A., Meuth, M., and Cedar, H. (1989) Mapping replication units in animal cells. Cell 57, 909–920

46. Hamlin, J. L., Dijkwel, P. A., and Vaughn, J. P. (1992) Initiation of replication in the Chinese hamster dihydrofolate reductase domain. Chromosoma 102, S17–23

47. Dijkwel, P. A., and Hamlin, J. L. (1992) Initiation of DNA replication in the dihydrofolate reductase locus is confined to the early S period in CHO cells synchronized with the plant amino acid mimosine. Molecular and cellular biology 12, 3715–3722

48. Kitsberg, D., Selig, S., Keshet, I., and Cedar, H. (1993) Replication structure of the human beta-globin gene domain. Nature 366, 588–590

49. Delidakis, C., and Kafatos, F. C. (1989) Amplification enhancers and replication origins in the autosomal chorion gene cluster of Drosophila. The EMBO journal 8, 891–901

50. Heck, M. M., and Spradling, A. C. (1990) Multiple replication origins are used during Drosophila chorion gene amplification. The Journal of cell biology 110, 903–914

51. Burhans, W. C., Vassilev, L. T., Caddle, M. S., Heintz, N. H., and DePamphilis, M. L. (1990) Identification of an origin of bidirectional DNA replication in mammalian chromosomes. Cell 62, 955–965

52. Little, R. D., Platt, T. H., and Schildkraut, C. L. (1993) Initiation and termination of DNA replication in human rRNA genes. Molecular and cellular biology 13, 6600–6613

53. Giacca, M., Zentilin, L., Norio, P., Diviacco, S., Dimitrova, D., Contreas, G., Biamonti, G., Perini, G., Weighardt, F., Riva, S., and et al. (1994) Fine mapping of a replication origin of human DNA. Proceedings of the National Academy of Sciences of the United States of America 91, 7119–7123

54. Vassilev, L., and Johnson, E. M. (1990) An initiation zone of chromosomal DNA replication located upstream of the c-myc gene in proliferating HeLa cells. Molecular and cellular biology 10, 4899–4904

55. Dijkwel, P. A., and Hamlin, J. L. (1995) The Chinese hamster dihydrofolate reductase origin consists of multiple potential nascent-strand start sites. Molecular and cellular biology 15, 3023–3031

56. de Cicco, D. V., and Spradling, A. C. (1984) Localization of a cis-acting element responsible for the developmentally regulated amplification of Drosophila chorion genes. Cell 38, 45–54

57. Levine, J., and Spradling, A. (1985) DNA sequence of a 3.8 kilobase pair region controlling Drosophila chorion gene amplification. Chromosoma 92, 136–142

58. Aladjem, M. I. (2004) The mammalian beta globin origin of DNA replication. Frontiers in bioscience: a journal and virtual library 9, 2540–2547

59. Austin, R. J., Orr-Weaver, T. L., and Bell, S. P. (1999) Drosophila ORC specifically binds to ACE3, an origin of DNA replication control element. Genes & development 13, 2639–2649

60. Kong, D., Coleman, T. R., and DePamphilis, M. L. (2003) Xenopus origin recognition complex (ORC) initiates DNA replication preferentially at sequences targeted by Schizosaccharomyces pombe ORC. The EMBO journal 22, 3441–3450

61. Kim, S. M., and Huberman, J. A. (1998) Multiple orientation-dependent, synergistically interacting, similar domains in the ribosomal DNA replication origin of the fission yeast, Schizosaccharomyces pombe. Molecular and cellular biology 18, 7294–7303

62. Lee, J. K., Moon, K. Y., Jiang, Y., and Hurwitz, J. (2001) The Schizosaccharomyces pombe origin recognition complex interacts with multiple AT-rich regions of the replication origin DNA by means of the AT-hook domains of the spOrc4 protein. Proceedings of the National Academy of Sciences of the United States of America 98, 13589–13594

63. Chuang, R. Y., Chretien, L., Dai, J., and Kelly, T. J. (2002) Purification and characterization of the Schizosaccharomyces pombe origin recognition complex: interaction with origin DNA and Cdc18 protein. The Journal of biological chemistry 277, 16920–16927

64. Le-Niculescu, H., Niesman, I., Fischer, T., DeVries, L., and Farquhar, M. G. (2005) Identification and characterization of GIV, a novel Galpha i/s-interacting protein found on COPI, endoplasmic reticulum-Golgi transport vesicles. J Biol Chem 280, 22012–22020

65. Anai, M., Shojima, N., Katagiri, H., Ogihara, T., Sakoda, H., Onishi, Y., Ono, H., Fujishiro, M., Fukushima, Y., Horike, N., Viana, A., Kikuchi, M., Noguchi, N., Takahashi, S., Takata, K., Oka, Y., Uchijima, Y., Kurihara, H., and Asano, T. (2005) A novel protein kinase B (PKB)/AKT-binding protein enhances PKB kinase activity and regulates DNA synthesis. J Biol Chem 280, 18525–18535

66. Kumagai, A., Shevchenko, A., and Dunphy, W. G. (2010) Treslin collaborates with TopBP1 in triggering the initiation of DNA replication. Cell 140, 349–359

67. Baltin, J., Leist, S., Odronitz, F., Wollscheid, H. P., Baack, M., Kapitza, T., Schaarschmidt, D., and Knippers, R. (2006) DNA replication in protein extracts from human cells requires ORC and Mcm proteins. J Biol Chem 281, 12428–12435

68. Berberich, S., Trivedi, A., Daniel, D. C., Johnson, E. M., and Leffak, M. (1995) In vitro replication of plasmids containing human c-myc DNA. J Mol Biol 245, 92–109

69. Krude, T., Jackman, M., Pines, J., and Laskey, R. A. (1997) Cyclin/Cdk-dependent initiation of DNA replication in a human cell-free system. Cell 88, 109–119

70. Vashee, S., Cvetic, C., Lu, W., Simancek, P., Kelly, T. J., and Walter, J. C. (2003) Sequence-independent DNA binding and replication initiation by the human origin recognition complex. Genes & development 17, 1894–1908

71. Lee, D. G., and Bell, S. P. (2000) ATPase switches controlling DNA replication initiation. Current opinion in cell biology 12, 280–285

